# Self-supervised learning enables robust microbiome predictions in data-limited and cross-cohort settings

**DOI:** 10.1101/2025.10.19.683269

**Authors:** Liron Zahavi, Zachary Levine, Anastasia Godneva, Veronika Dubinkina, Raja Dhir, Katherine S. Pollard, Adina Weinberger, Eran Segal

## Abstract

The gut microbiome plays a crucial role in human health, but machine learning applications in this field face significant challenges, including limited labeled data availability, high dimensionality, and batch effects across different cohorts. To address these limitations, we developed representation learning models for gut microbiome metagenomic data, drawing inspiration from foundation models approaches based on self-supervised and transfer learning principles. By leveraging a large collection of 85,364 metagenomic samples, we implemented multiple self-supervised learning methods, including masked autoencoders with varying masking rates and adapted single-cell RNA sequencing models (scVI and scGPT), to generate embeddings from bacterial abundance profiles. These learned representations demonstrated significant advantages over raw bacterial abundances in two key scenarios: first, when training predictive models with very limited labeled data, improving prediction performance for age (r = 0.14 vs. 0.06), BMI (r = 0.16 vs. 0.11), visceral fat mass (r = 0.25 vs. 0.18), and drug usage classification (PR-AUC = 0.81 vs. 0.73); and second, when generalizing predictions across different cohorts, consistently outperforming models based on raw abundances in cross-dataset evaluation. Our approach provides a valuable framework for leveraging self-supervised representation learning to overcome the data limitations inherent in microbiome research, potentially enabling more robust and generalizable machine learning applications in this field.

## Introduction

The gut microbiome exerts widespread influence on human health through its interconnected effects on metabolism, immunity, and neurological functions. This dynamic community of microorganisms operates through complex ecological networks and host interactions, which remain only partially understood. Analysis of metagenomic data from gut microbiome samples offers valuable insights into these complex microbial communities.

A significant challenge in microbiome research is the limited availability of data. Shotgun metagenomic sequencing remains costly, and studies often include relatively small cohorts. This creates a fundamental obstacle for machine learning applications, as the data is inherently high-dimensional, typically represented as relative abundances across thousands of microbial species and millions of genes. The combination of high dimensionality with small sample sizes leads to models that struggle with generalizability and robustness.

A natural solution to limited sample sizes would be to combine multiple cohorts, but this approach introduces its own challenges. Batch effects in metagenomic data make it difficult to combine multiple cohorts or compare different populations. These technical variations can mask true biological signals, limiting our ability to leverage diverse datasets to increase effective sample size.

Self-supervised and transfer learning approaches have shown remarkable success in other biological domains, including protein structure prediction, genomic analysis, and single-cell transcriptomics ^1 2 3 4^. These deep-learning methods leverage large amounts of unlabeled samples to learn the structure of the data, training with labels that are generated from the data itself. By incorporating knowledge learned from the large datasets, they can be used to create meaningful sample representations that transfer well to downstream tasks with limited labeled data. Importantly, these approaches can mitigate overfitting to dataset-specific signals, including batch effects, by learning more generalizable patterns across diverse data sources.

These principles are particularly relevant for biological applications where labeled data is scarce but unlabeled data is abundant. They enable the processing of samples from new cohorts with limited sample sizes, leveraging patterns learned from larger datasets to extract meaningful features. In the microbiome research field, large datasets are starting to accumulate — but they typically include very minimal host information, which limits supervised learning applications. While recent work has begun exploring microbiome representation learning ^5 6 7 8^, mainly using 16S sequencing data (which offers lower resolution than metagenomics), this emerging field still offers significant opportunities for methodological advancement. Critically, many representation learning works in biology fail to demonstrate clear advantages over standard approaches used in the field, leaving their practical utility unclear ^9,10^.

To address these challenges, we developed representation learning models for gut microbiome shotgun metagenomics that leverage large-scale unlabeled datasets totalling 85,364 samples. We explored multiple architectural approaches, including masked autoencoders and cross-domain adaptation of established single-cell RNA sequencing models. Our study makes three key methodological contributions: (1) benchmarking that demonstrates clear advantages over the standard practice in microbiome machine learning --- prediction from raw bacterial relative abundances; (2) novel cross-domain transfer of successful single-cell RNA sequencing models to microbiome analysis; and (3) a pretrained model that can be used to extract representations from new metagenomic samples.

Our learned representations demonstrate clear advantages over standard microbiome analysis approaches in two critical scenarios: enhanced performance in limited-data settings that reflect real-world study constraints, and improved generalizability across different populations and technical frameworks.

These advances directly address critical barriers to machine learning applications in microbiome research and establish a framework for adapting successful biological representation learning methods across related domains.

### Representation learning models

To create meaningful representations of gut microbiome samples, we employed self-supervised learning (SSL), in which deep learning models learn from unlabeled data by creating learning objectives directly from the data structure itself. We gathered metagenomic data from three studies: a cohort of 66,151 participants from both the US (6,206; “1000_US”) and Israel (59,945; “1000_IL”) ^11^, a cohort of 11,084 participants from Israel (“10_IL”) ^12,13^, and a cohort of 8,129 participants from the Netherlands (“32_NE”) ^14^. These datasets span different geographies, ancestries, collection periods and laboratories, providing a robust foundation for learning generalizable microbiome representations.

All datasets included metagenomic sequencing data alongside basic host information (age, sex, Body Mass Index (BMI)), with one cohort (10_IL, ^13^) also providing diverse clinical and phenotypic measurements, including body composition profile from medical imaging, drug usage patterns, and disease diagnoses. Our approach involved a two-stage process: first, we used only metagenomic data to train the models and learn representations. Subsequently, we evaluated the effectiveness of these learned representations by testing their impact on machine learning models predicting host phenotypes.

For each sample, we mapped sequencing reads to a reference database of bacterial genomes and retained the 902 most prevalent species after filtering (requiring species to be present in at least 500 samples of the deeply phenotyped cohort; Methods). We calculated relative species abundances, creating a species abundance vector (902 dimensions) for each sample — a standard representation of microbiome samples. This vector serves as both our baseline representation (“raw’’, Table 1) and the input to our representation learning models.

**Table 1.** Prediction performance across different microbiome representations. Performance evaluation was based on four repetitions of a 5-fold cross-validation, resulting in a total of 20 train-test split experiments. The table shows the performance (continuous: Pearson correlation, binary: PR-AUC) mean and standard deviation across all test sets. Q-values are based on the Mann-Whitney U test comparing each performance distribution to the raw model’s performances, and corrected for multiple testing using FDR.

|  | raw | scVI | scGPT | mask-AE(10%) | mask-AE(30%) | mask-AE(70%) | mask-transformer (30%)-mean & max |
| --- | --- | --- | --- | --- | --- | --- | --- |
| age, mean (std) | 0.34 (0.02) | 0.36 (0.01) | 0.32 (0.02) | 0.34 (0.01) | **0.37 (0.01)** | 0.36 (0.01) | 0.33 (0.02) |
| age, q-value |  | q=0.05 | n.s. | n.s. | q=0.0003 | q=0.002 | n.s. |
| gender, mean (std) | **0.67 (0.02)** | 0.66 (0.02) | 0.66 (0.01) | **0.67 (0.01)** | **0.67 (0.01)** | 0.66 (0.01) | 0.66 (0.01) |
| gender, q-value |  | n.s. | n.s. | n.s. | n.s. | n.s. | n.s. |
| bmi, mean (std) | 0.30 (0.02) | 0.30 (0.02) | 0.29 (0.02) | 0.30 (0.02) | **0.31 (0.02)** | 0.30 (0.02) | 0.28 (0.02) |
| bmi, q-value |  | n.s. | n.s. | n.s. | n.s. | n.s. | n.s. |
| total_scan_vat_mass, mean (std) | 0.37 (0.02) | 0.37 (0.02) | 0.37 (0.02) | 0.37 (0.02) | **0.38 (0.02)** | **0.38 (0.02)** | 0.35 (0.02) |
| total_scan_vat_mass, q-value |  | n.s. | n.s. | n.s. | q=0.05 | q=0.09 | n.s. |
| is_PPI, mean (std) | **0.21 (0.04)** | 0.18 (0.05) | 0.08 (0.02) | **0.21 (0.04)** | **0.21 (0.04)** | **0.21 (0.04)** | 0.09 (0.02) |
| is_PPI, q-value |  | n.s. | n.s. | n.s. | n.s. | n.s. | n.s. |
| is_Hyperlipidemia, mean (std) | 0.24 (0.02) | 0.23 (0.01) | 0.24 (0.02) | 0.24 (0.01) | **0.25 (0.01)** | 0.24 (0.02) | 0.24 (0.01) |
| is_Hyperlipidemia, q-value |  | n.s. | n.s. | n.s. | q=0.4 | q=0.7 | n.s. |
| is_FattyLiverDisease, mean (std) | **0.09 (0.02)** | 0.08 (0.02) | **0.09 (0.02)** | **0.09 (0.02)** | **0.09 (0.02)** | 0.08 (0.01) | 0.08 (0.01) |
| is_FattyLiverDisease, q-value |  | n.s. | q=0.6 | n.s. | n.s. | n.s. | n.s. |

To learn sample representations, we implemented multiple SSL approaches. First, we trained a masked autoencoder. Masking is a common SSL approach, in which a proportion of the sample inputs are obscured and the model is trained to reconstruct them. This approach is designed to learn underlying microbial community patterns by forcing the model to predict missing species based on observed ones, with the expectation of capturing ecological relationships and functional redundancies within microbial communities. We masked the abundances of 10, 30, or 70% of the species in each sample, testing low, medium, and high masking fractions. Starting with a basic deep-learning architecture, we trained multi-layer perceptron (MLP) models to reconstruct the hidden values, and used the encoded representation from the middle layer as the sample representation. We set aside 10% of the data as a validation set, and used it to choose a stopping point for training. In the self-supervised task of reconstructing masked abundances, the 10%, 30%, and 70% masking models achieved an explained variance (EV) of 26.4%, 24.5%, and 19.1%, respectively, in the validation set averaged across all species *(Supp. figs. 1-3)*.

We also evaluated a transformer-based masked autoencoder. Transformers, originally developed for natural language processing, use self-attention mechanisms that can capture complex dependencies between all input features simultaneously. This architecture offered the potential to model ecological interactions between bacterial species better than the simpler feedforward networks. In our implementation, we stratified the relative abundances of each species into quantiles, assigned a token per species quantile, and applied the same masking-reconstruction task as with the MLP models. To extract sample representations for downstream tasks, we mean-and max-pooled the output of the trained model (Methods).

Additionally, recognizing the shared characteristics between metagenomic and single-cell RNA sequencing (scRNA-seq) data – both involve high-dimensional, sparse (zero-inflated), tabular data, with features (genes or species) exhibiting highly variable abundance distribution, and pronounced batch effects – we explored the potential of adapting well-established representation learning models from the scRNA-seq domain. While representation learning for microbiome data remains in its nascent stages, the scRNA-seq field has advanced significantly in recent years, with numerous foundation models designed specifically to handle high-dimensional, sparse biological data. We hypothesized that the underlying principles of these models, designed to capture complex biological variations and mitigate technical noise in scRNA-seq data, could be effectively transferred to the analysis of gut microbiome compositions. To this end, we adapted and evaluated two prominent scRNA-seq representation learning models: scVI ^3^ and scGPT ^15^.

scVI (single-cell Variational Inference) is a variational autoencoder (VAE)-based model that learns a low-dimensional latent space by modeling observed data as arising from a probabilistic generative process. In our adaptation, we treated bacterial species abundances analogously to gene expression levels, allowing the model to capture the complex distribution of microbiome compositions. This approach enables explicit incorporation of batch information, helping to disentangle technical variation from biological signals. The probabilistic framework of scVI is particularly well-suited for addressing zero-inflation, inherent measurement noise, and the varying abundance distributions across different bacterial species that characterize both scRNA-seq and metagenomic data, potentially yielding more robust representations of microbial communities.

Conversely, scGPT represents a foundation model approach for single-cell omics data, utilizing transformer architectures to capture complex patterns in sparse, high-dimensional biological data. Its self-attention mechanisms, originally designed to model gene-gene interactions in single cells, could potentially capture ecological relationships between bacterial species in microbiome communities. We adapted scGPT for microbiome analysis, treating bacterial species abundances analogously to gene expression values, with the hypothesis that its specialized architecture for modeling biological interactions would effectively capture patterns of bacterial co-occurrence and mutual exclusion in gut microbiome samples.

In addition to species abundance information, we provided the scVI and scGPT models with batch and cohort information, demonstrating a multi-modal approach that jointly models biological features with technical covariates to handle technical variation and biological heterogeneity.

### Sample embeddings improve phenotype predictions from microbiome

We designed comprehensive evaluations to rigorously test whether our learned representations provide advantages over the current standard practice in microbiome machine learning: using raw bacterial relative abundances as features for predictive models. We leveraged a dataset with rich host information to train tree-based models ^16^ to predict host phenotypes, comparing the performance of models trained on sample embeddings to those trained on raw bacterial relative abundances — the standard approach in microbiome machine learning. We performed 4 repetitions of 5-fold cross-validation to robustly assess the predictive performance of different representations. We trained models to predict host age, sex, BMI, Visceral Adipose Tissue (VAT) mass, Proton Pump Inhibitors (PPI) intake, hyperlipidemia diagnosis, and fatty liver disease diagnosis.

For the first benchmarking analysis, we used the entire deeply phenotyped cohort (n = 11,084; Table 1). Age prediction based on the 30 and 70% mask-AE models embedding was better than the prediction from the raw relative abundances, with mean Pearson correlations of 0.34 for the raw features, 0.37 for the 30% mask-AE (q-value, FDR adjusted = 0.0003), and 0.36 for the 70% mask-AE (q-value = 0.002). The superiority of these models’ embeddings in predicting Visceral Adipose Tissue (VAT) mass was marginally significant, with a Pearson correlation of 0.38 for both these models, and 0.37 for the raw abundances (q-value = 0.05, 0.09). In other cases, we did not observe a significant advantage for models trained on the learned representations rather than the bacterial abundances. For the transformer model, the different pooling strategies we tested yielded similar results (Supp. table 1).

In the second benchmarking analysis, we tested the benefit of the learned representations in limited-data scenarios. We hypothesized that in scenarios in which there are much fewer labels to learn from, learning features in an SSL scheme will be more advantageous, as small cohorts lack the diversity and sample size needed to learn complex microbiome relationships from scratch. To simulate this, we downsampled our cohort to 100 training samples — a typical scale for many microbiome studies, clinical trials in particular — and used them to train the phenotype predictors from either the representations or the raw bacterial relative abundances (Table 2, Supp. table 2).

**Table 2.** Prediction performance across microbiome representations with limited (n=100) training data. Train cohorts consisted of 100 randomly selected samples. Performance distribution is based on 100 repetitions. The table shows the performance (continuous: Pearson correlation, binary: PR-AUC) mean and standard deviation across all test sets. Q-values are based on the Mann-Whitney U test comparing each performance distribution to the raw model’s performances, and corrected for multiple testing using FDR.

|  | raw | scVI | scGPT | mask-AE(10%) | mask-AE(30%) | mask-AE(70%) | mask-transformer (30%)-mean & max |
| --- | --- | --- | --- | --- | --- | --- | --- |
| age, mean (std) | 0.06 (0.11) | **0.14 (0.11)** | 0.13 (0.09) | 0.10 (0.11) | 0.11 (0.11) | 0.11 (0.12) | 0.12 (0.10) |
| age, q-value |  | q<10<sup>-5</sup> | q<10<sup>-4</sup> | q=0.08 | q=0.003 | q=0.004 | q=0.002 |
| gender, mean (std) | 0.62 (0.07) | 0.60 (0.07) | 0.63 (0.06) | 0.62 (0.06) | **0.64 (0.06)** | **0.64 (0.06)** | 0.61 (0.06) |
| gender, q-value |  | n.s. | q=0.2 | q=0.7 | q=0.2 | q=0.08 | n.s. |
| bmi, mean (std) | 0.11 (0.11) | 0.12 (0.11) | 0.15 (0.09) | 0.15 (0.11) | **0.16 (0.11)** | **0.16 (0.11)** | 0.12 (0.11) |
| bmi, q-value |  | q=0.8 | q=0.03 | q=0.08 | q=0.003 | q=0.02 | q=0.7 |
| total_scan_vat_mass, mean (std) | 0.18 (0.11) | 0.18 (0.10) | 0.20 (0.10) | 0.23 (0.10) | **0.25 (0.11)** | 0.24 (0.11) | 0.18 (0.11) |
| total_scan_vat_mass, q-value |  | q=0.9 | q=0.4 | q=0.004 | q=0.0004 | q=0.003 | q=0.8 |
| is_PPI, mean (std) | 0.73 (0.08) | 0.76 (0.06) | 0.61 (0.06) | 0.79 (0.06) | 0.80 (0.05) | **0.81 (0.04)** | 0.62 (0.06) |
| is_PPI, q-value |  | q=0.009 | n.s. | q<10<sup>-5</sup> | q<10<sup>-9</sup> | q<10<sup>-11</sup> | n.s. |
| is_Hyperlipidemia, mean (std) | **0.53 (0.05)** | **0.53 (0.05)** | **0.53 (0.05)** | 0.52 (0.06) | 0.52 (0.05) | 0.52 (0.05) | **0.53 (0.05)** |
| is_Hyperlipidemia, q-value |  | q=0.6 | q=0.8 | q=0.9 | q=0.9 | q=0.8 | q=0.7 |
| is_Fatty Liver Disease, mean (std) | 0.57 (0.05) | 0.55 (0.06) | 0.58 (0.06) | 0.58 (0.06) | **0.59 (0.07)** | 0.58 (0.06) | 0.57 (0.06) |
| is_Fatty Liver Disease, q-value |  | n.s. | q=0.3 | q=0.4 | q=0.1 | q=0.5 | q=0.8 |

Indeed, the advantage of representation learning became much more pronounced in this data-limited setting. Predicting age from the learned representations improved the performance from a Pearson correlation of 0.06 to 0.14 (scVI, q-value<10^-5^). The Pearson correlation for BMI prediction improved from 0.11 to 0.16 (mask-AE 30%, q-value=0.003), and VAT mass prediction improved from 0.18 to 0.25 (mask-AE 30%, p-value=0.0004). In the classification of individuals using proton pump inhibitor (PPI) drugs, PR-AUC improved from 0.73 to 0.81 (mask-AE 70%, p-value<10^-11^). In the classification of sex and fatty liver disease, the mean PR-AUC achieved with the learned representations was also better than that gained with the raw abundances, but the difference was not statistically significant. This demonstrates the advantage of learned representations from SSL models in training supervised models on small datasets.

### Microbiome predictors trained on learned representations are more generalizable to new cohorts

Microbiome datasets differ due to both biological and technical factors. Sample collection and storage, DNA extraction, PCR amplification and library preparation, sequencing platforms and runs, and data processing protocols all affect the observed species composition. In addition, environmental variables and health differences between populations are reflected in different microbiome profiles. These differences impair the ability to apply machine learning models trained on one dataset to label samples from another dataset — a critical requirement for clinical deployment. We hypothesized that using features learned by the SSL models, rather than the raw bacterial abundances, would enable models to generalize better between datasets.

To test if the learned representations improve the generalizability of supervised models, we separated our dataset to four subsets (by study and country), trained tree-based models predicting age, sex, and BMI on each subset, and evaluated it on the other three (Supp. fig. 4; Methods). Notably, we did not finetune the supervised models on the target cohort — only predicted the labels of the target cohort using the models that were fitted on the source cohort, testing how well the models generalize across different populations. Overall, across five representation models and three phenotypes, the learned representations improved cross-cohort prediction in 148 of 180 comparisons (82%) relative to raw features, with an average improvement of 10% ± 15% (Fig. 1, Supp. Fig. 4). When transferring age and BMI models to different cohorts, scVI and the three mask-AE representations each outperformed the raw-based model in 10 to 12 of the 12 train-test cohort pairs (q-values=0.03 to 0.001). The highest improvement was demonstrated when applying the age prediction model that was trained on the 32_NE cohort to predict age in the 10_IL cohort — the model based on mask-AE 30% representations outperformed the raw-based model by 81%. For sex classification, improvements were more modest, achieving an increase in PR-AUC of up to 14% (q-values=0.09 to 0.007). The only model whose representations did not significantly outperform the baseline was scGPT. These results demonstrate the advantage of learned microbiome representations as features for models that generalize better across cohorts.

**Figure 1.**
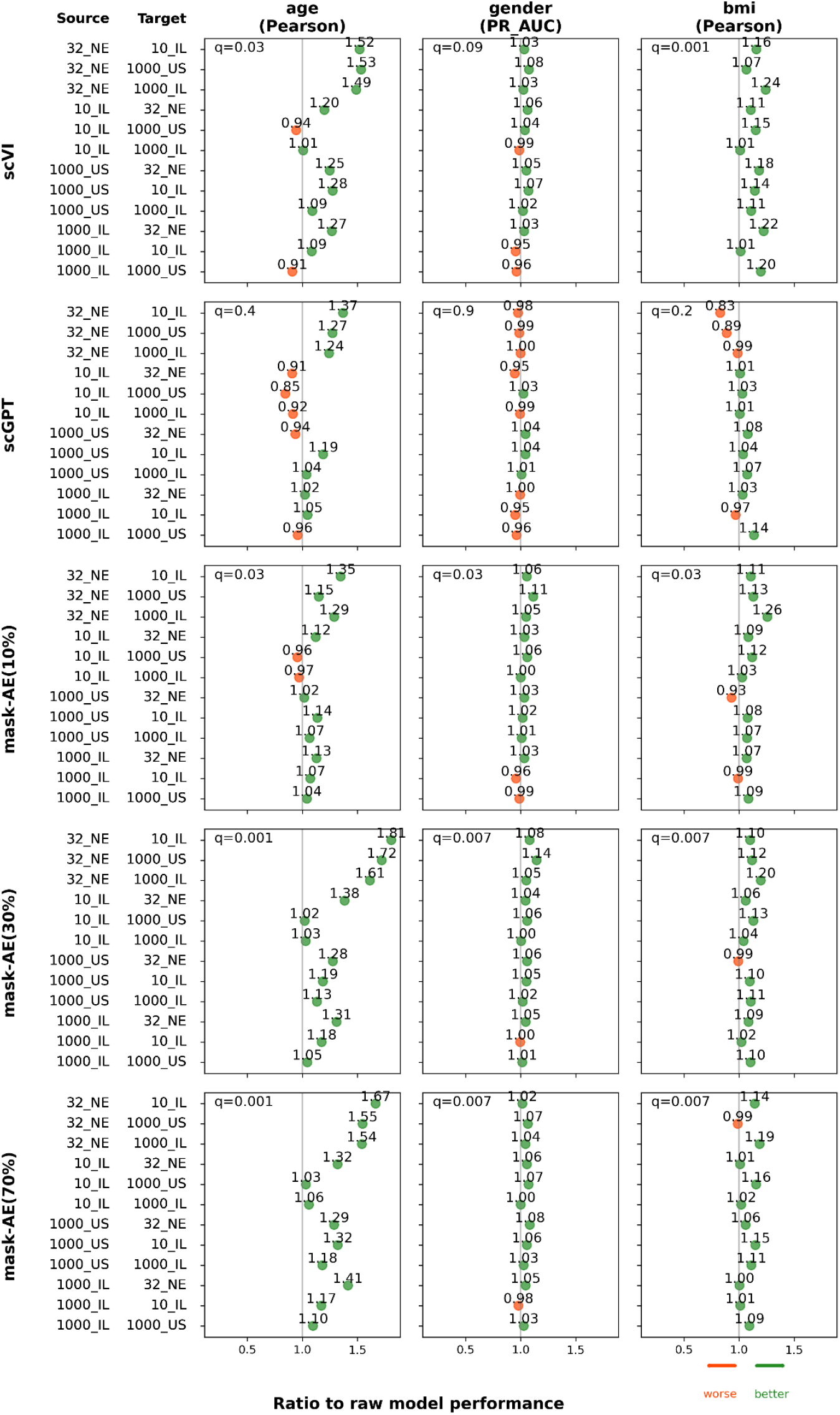
cross-cohort generalizability of models compared to the baseline model. Each point represents a train-test cohort pair, showing the ratio of representation-based model performance to raw abundance model performance. Values >1.0 indicate superior performance of learned representations. Q-values are FDR-adjusted. Original performance metrics of each model are detailed in Supp. Fig. 4.

### Model application to new studies

To facilitate reproducibility and enable broader application of our findings, we make our best-performing masked autoencoder model available to the microbiome research community. The pretrained 30% masked autoencoder model, which demonstrated superior performance in our evaluations, is available along with example code showing how to generate embeddings from bacterial abundance profiles.

The pretrained model accepts the relative abundances of bacterial species in samples and outputs the corresponding embedding vectors that can be used for downstream machine learning tasks. These embeddings may be particularly valuable for improving performance in scenarios with limited training data or when attempting to develop models that generalize across different cohorts.

## Discussions

In this study, we developed representation learning models for gut microbiome metagenomic data and demonstrated that self-supervised learning can significantly improve microbiome-based predictive modeling in two critical scenarios: limited-data settings and cross-cohort generalization. These improvements address fundamental challenges in microbiome machine learning with clear implications for clinical translation.

Self-supervised learning and transfer learning approaches have recently emerged in microbiome research ^5 6 7 8^. Contributing to this emerging area, our work leverages a large metagenomic dataset of 85,364 samples to train multiple representation learning models and systematically evaluates their performance against the current standard practice in microbiome machine learning — using raw bacterial abundances with standard machine learning methods — across diverse prediction scenarios, addressing a fundamental need to demonstrate practical utility over established methods ^9,10^.

Particularly noteworthy was the consistent superiority of representation-based models in cross-cohort prediction tasks, improving prediction accuracy by 10% on average and up to 81% compared to models trained on raw features. The ability of our models to generalize across different populations and data collection protocols addresses one of the most significant challenges in microbiome machine learning: batch effects and population-specific signatures that typically impair model transferability. By learning more robust, generalizable features that capture underlying biological signals rather than technical variations, our approach offers a promising avenue for developing more universal microbiome-based predictive models. This enhanced generalizability addresses one of the most significant barriers to clinical translation of microbiome research: the need for models that perform reliably across diverse populations, healthcare systems, and technical protocols.

The significant improvements in data-limited settings demonstrate how self-supervised pretraining enables effective knowledge transfer, allowing models to leverage patterns learned from large unlabeled datasets when supervised training data is scarce. While the performance gains were modest when using the full dataset, they became substantially more pronounced when restricting training to just 100 samples. This pattern aligns with findings in other domains where representation learning shows the greatest advantage in low-data regimes, effectively leveraging patterns learned from larger unlabeled datasets to improve performance on specific tasks with limited labeled examples, and is particularly relevant for microbiome research. While our deeply-phenotyped cohort contains over 10,000 samples, the majority of microbiome studies operate with cohorts 1-3 orders of magnitude smaller, typically consisting of tens to hundreds of samples. These smaller studies often focus on specific conditions or populations that cannot feasibly be expanded to larger scales. This capability could enable meaningful discoveries in studies that were previously underpowered due to sample size constraints.

Among the representation models we evaluated, the 30% masked autoencoder consistently demonstrated the best overall performance across prediction tasks, followed by the 70% masking model. This suggests that intermediate levels of feature masking may provide an optimal balance between preserving sufficient information for meaningful reconstruction while forcing the model to learn robust underlying patterns. Despite the theoretical advantages of transformer architectures in capturing complex inter-species relationships through self-attention mechanisms, our transformer-based model did not outperform the simpler MLP-based approaches. This pattern was mirrored in the adapted single-cell models: while scVI showed competitive performance and even outperformed other methods in specific scenarios (age prediction in small cohorts and BMI prediction in cross-cohort analyses), the more sophisticated transformer-based scGPT model generally underperformed relative to both scVI and our custom models.

Our work builds upon recent successes of representation learning in other biological domains, particularly in single-cell RNA sequencing. The scRNA-seq field has developed sophisticated computational approaches to overcome challenges that are similar to those in microbiome research: high dimensionality, technical variation, zero-inflated sparse data, and limited sample sizes. Models like scVI have become standard tools for integrating and analyzing diverse scRNA-seq datasets, suggesting similar approaches could benefit microbiome research. Our successful adaptation of single-cell RNA sequencing model architectures to microbiome analysis demonstrates the value of cross-domain methodological transfer in biological data analysis, leveraging mature computational methods from one biological domain to advance another.

These advances have practical implications for microbiome research applications. By enabling smaller studies to achieve meaningful predictive performance and improving model transferability across populations, our approach could accelerate the identification of diagnostic microbiome signatures and support the development of models that perform consistently across diverse cohorts. This enhanced analytical capability may be particularly valuable for studies involving rare conditions, controlled interventions, or unique environmental exposures, where large sample collection is challenging. To support these applications, we provide our best-performing pretrained model for community use.

While our training dataset is large by microbiome standards, it remains modest compared to foundation model datasets in other domains. Future work could benefit from even larger datasets, representing a bigger variety of geographies and health states. Other models built upon this framework and an even larger dataset may include a larger selection of species — including the rare ones — as well as greater resolution of features, like strains and genes. Our representations also sacrifice some interpretability compared to raw abundance approaches, though the performance gains may justify this trade-off in many applications.

In conclusion, we demonstrate that self-supervised learning on large metagenomic datasets can substantially enhance model generalizability across diverse populations and improve prediction accuracy in data-limited scenarios. This work establishes representation learning as a viable solution for overcoming fundamental challenges in microbiome-based machine learning, opening new possibilities for leveraging larger unlabeled datasets to improve predictions in specialized studies where large labeled cohorts remain challenging to obtain.

## Methods

### Data

We based our work on data from three previously published studies: a cohort of 66,151 individuals from the US and from Israel (“1000_US” and “1000_IL”, accordingly) ^11^, a cohort of 8,129 individuals from the Netherlands (“32_NE”) ^14^, and a cohort of 11,084 individuals from Israel (“10_IL”) ^12,13^. All datasets included gut microbiome metagenomic data alongside basic host information: age, sex, and BMI. From the 10_IL cohort, we also obtained data on body composition, drug usage, and medical diagnoses. We only included the earliest sample of each participant.

### Preprocessing the microbiome data

We processed the metagenomic samples and aligned them to bacterial reference genomes as described in Leviatan et al. ^17^ to infer species composition. We filtered the species by prevalence, including species seen in at least 500 samples in the deeply-phenotyped (“10_IL”) cohort (out of 11,084 samples) — resulting in 902 species. For these 902 species, we calculated their relative abundances in each sample (summing to 1). Our pipeline’s detection threshold is set to 0.0001, and we truncated any smaller abundance value to this minimum.

### Masked autoencoder

To create sample representations using masked autoencoders, we implemented multi-layer perceptron (MLP) architectures trained on a self-supervised reconstruction task. We preprocessed bacterial relative abundance data by applying a log10 transformation and shifting values by 2 to convert the range from [-4, 0] to [-2, 2], facilitating more stable neural network training.

We employed a masking scheme where we randomly obscured 10%, 30%, or 70% of input features for each sample during training, replacing masked positions with a fixed value of -2. The masking pattern was randomly generated for each sample in every training batch. We computed the mean squared error loss only on the obscured positions.

We optimized hyperparameters through a systematic hyperparameter sweep, testing combinations of masking fractions, encoding dimensions, number of layers, learning rates, batch sizes, dropout rates, activation functions, masking values, whether to shift values by 2 or not, and loss computation strategies (loss on all features vs. masked features only). For each masking fraction, we selected the best-performing hyperparameter combination, resulting in models with different optimal configurations. The 10% and 70% masking models employed Mish activation with 10% dropout, while the 30% masking model used GELU activation with 25% dropout. All final models used a learning rate of 0.0005 with Adam optimization, training for 300-350 epochs. Our autoencoder architecture consisted of an encoder-decoder structure with single-layer components and batch normalization. The encoder transformed the 902-dimensional input (representing bacterial species abundances) to a 1024-dimensional latent representation, while the decoder reconstructed the original input dimensions.

We split the dataset into 90% training and 10% validation sets, monitoring reconstruction performance using mean squared error loss and explained variance. The learned representations were extracted from the encoder’s output layer after training was completed.

### Masked transformer

We also implemented a transformer-based masked autoencoder using self-attention mechanisms to capture potential interactions between bacterial species. For input representation, we tokenized bacterial abundance values using species-specific quantile binning. For each bacterial species independently, we divided the abundance range into 10 bins: one bin for the minimal value (-4), and 9 bins based on quantiles of non-minimal abundances. Each species-abundance combination was assigned a unique token, creating a vocabulary of 9,021 tokens (902 species × 10 bins + 1 mask token). This quantile-based approach preserves relative abundance information while creating discrete tokens suitable for transformer processing.

The transformer architecture consisted of 2 encoder layers with 256-dimensional embeddings, 16 attention heads, and feedforward layers with 512 hidden units. We employed a 10% dropout and trained the model as a regression task, predicting continuous abundance values rather than discrete tokens. Each bacterial species had its own linear decoder head that converted the transformer’s output embeddings to abundance predictions.

We applied the same masking strategy as the MLP models, randomly masking 30% of input features and replacing them with a special mask token. The mean squared error loss was computed only on the masked and reconstructed inputs. We used Adam optimization with a learning rate of 0.001, a batch size of 256, and trained for 260 epochs with a 90%-10% train-validation split. The learned representations were extracted by max-pooling, mean-pooling, or by concatenating mean-pooling and max-pooling of the transformer’s output across the species dimension, resulting in 512-dimensional sample representations (256 dimensions each from mean and max pooling).

### scVI

To create sample representations with scVI ^3^, we used scVI-tools version 1.2.2.post2 with Python 3.11.5. To align with the expected input format of the scRNA-seq models, which are typically applied to count data, we converted bacterial relative abundances to pseudo-counts by multiplying each sample’s relative abundances by its total mapped read count, then converted the resulting values to integers. We replaced abundance values of 0.0001 with zero to reflect the detection limit of our data.

We constructed batch identifiers combining country, study type, DNA extraction kit, library preparation method, and collection year to account for technical variation across our diverse datasets. We configured scVI with a 2-layer encoder-decoder architecture using 512 hidden units and 256 latent dimensions. To model the count distribution, we used the zero-inflated negative binomial (ZINB) distribution with gene-batch-specific dispersion parameters — to model the sparsity and the variability of distributions between species and batches that is common in microbiome data. We enabled covariate encoding and used embedding-based batch representation with 5-dimensional batch embeddings.

We trained the model from scratch for up to 50 epochs using a batch size of 128 and a validation set comprising 5% of samples. We implemented early stopping with a patience of 3 epochs and used a learning rate of 0.001 with KL divergence warmup over 6 epochs. Learning rate reduction on plateau was enabled with a patience of 2 epochs. After training, we extracted the latent representations from the trained model to serve as sample embeddings for downstream analysis.

### scGPT

To create sample representations with scGPT ^15^, we used scGPT 0.2.1 with Python 3.11.3 and Torch 2.3.1. We trained on each sample’s vector of species abundance, tokenizing each species abundance so that a sequence of abundance values became a sequence of tokens. We used a masking ratio of 0.4 and 25 bins to tokenize the relative abundance values. As we did not have a cell type feature, we disabled the elastic cell similarity objective by setting its relative weight to zero. We used the sequencing run batch label (“RunName”) as the batch identifier for batch integration.

We chose the transformer hyperparameters based on the lowest masked MSE evaluation loss (L2 norm between true and predicted values for masked tokens), and tested embedding dimensions of 128 to 768, and depths between two to six. We trained the final model from scratch using a learning rate of 1e-3, a batch size of 64, and set the number of transformer encoder layers to three, using 32 heads and a hidden token dimension of 256. Automatic mixed precision was used. We fixed the random seed for the training experiment to 42. We used the masking value of -1. We set abundance values of 0.0001 to zero in keeping with the detection limit, and ignored ‘cell type’ as it was not relevant for our use case. We created a vocabulary based on species names as opposed to gene names. We used the data in log1p form. We trained for 10 epochs, using all other default settings. We extracted the CLS token (cell embeddings in the original paper) after training as the sample representation for downstream analysis.

### Full cohort phenotype predictions

We conducted an analysis to evaluate the advantage of the various models for generating features for supervised models. For this analysis, we used the data from the 10_IL cohort, which includes information about host health. We trained tree-based models to predict host age, sex, Body Mass Index (BMI), Visceral Adipose Tissue (VAT) mass, Proton Pump Inhibitors (PPI) intake, hyperlipidemia diagnosis, and fatty liver disease diagnosis, from either the raw microbiome abundances, or the sample embeddings extracted from the pretrained models. It is important to note that the embeddings were extracted from the self-supervised models, which were trained without the sample labels. We used the LightGBM package ^18^ with 2000 estimators, a learning rate of 0.001, max_depth of 3, min_child_samples of 30, 70% samples sampling, 60% features sampling, and an initial random seed of 42. We used a five-fold, stratified (for binary phenotypes), cross-validation scheme, and used 15% of the train samples as a validation set to enforce early stopping (after 80 rounds without improvement). We evaluated model performances on the test set using Pearson’s correlation for continuous phenotypes and the area under the precision recall curve (PR-AUC) for binary phenotypes. We conducted four repetitions of these experiments, for a total of 20 sets of train, validation, and test samples (over five folds and four repetitions) — from which we derived the mean and the standard deviation. To compare the test prediction performances of each model, we compared this 20-long vector with that of the predictor trained on species relative abundances using the Mann-Whitney U test ^19^ -(alternative=‘greater’). We adjusted for multiple testing across all model-phenotype comparisons using the Benjamini-Hochberg False Discovery Rate (FDR) procedure ^20^.

### Downsampling analysis

To evaluate prediction performances of models trained on 100 labeled samples, we conducted an experiment similar to the one described in the previous paragraph, with a few changes. For each model, we randomly selected 100 samples for the training set and 100 samples for the test set. For the binary phenotypes, we sampled 50 samples from each of the label groups. In this experiment, we did not apply early stopping, but reduced the number of estimators to 1000. We also did not use cross validation. For each pretrained model we compared and each target phenotype, we repeated this experiment 100 times, and used the Mann-Whitney U test to compare the performances to those gained by the model trained on species relative abundances. We adjusted for multiple testing across all model-phenotype comparisons using Benjamini-Hochberg False Discovery Rate (FDR) procedure ^20^.

### Cross-cohort predictions analysis

To test how well models trained on different representations predict phenotypes in cohorts different from those used to train them, we each time trained a prediction model on one cohort, and tested its prediction performance in other cohorts. To train the models, we used 85% of the cohort samples, and used the remaining 15% both for the within-cohort evaluation, and for performing early stopping (after not improving for 50 rounds). Then, to evaluate the trained model on a different target cohort, we tested it on 100% of the target cohort samples. For this reason, in this analysis we only performed a single evaluation per cohort pair, and did not have a distribution of performance values over folds and repetitions like in the previous analyses. For the supervised models we used LightGBM, with 2000 estimators and the same parameters used in the previous analyses. We adjusted for multiple testing across all model-phenotype comparisons using Benjamini-Hochberg False Discovery Rate (FDR) procedure.

## Model availability

The best performing trained model, MAE-30%, is available at https://github.com/LironZa/MBEmbed along with code to create representations of new metagenomic samples for machine learning applications with new microbiome datasets.

## Author contribution

L.Z. conceived the project, designed and conducted all analyses, interpreted the results, and wrote the manuscript. Z.L. performed data analysis and reviewed the manuscript. A.G. performed data analysis. V.D., R.D., K.S.P. and A.W. reviewed the manuscript. E.S. conceived, directed and designed the project and analyses, interpreted the results and wrote the manuscript.

## Acknowledgements

We thank the members of the Segal group for fruitful discussions. E.S. is supported by the Crown Human Genome Center; the Larson Charitable Foundation New Scientist Fund; the Else Kröner Fresenius Foundation; the White Rose International Foundation; the Ben B. and Joyce E. Eisenberg Foundation; the Nissenbaum Family; Marcos Pinheiro de Andrade and Vanessa Buchheim; Lady Michelle Michels; Aliza Moussaieff; and grants funded by the Minerva Foundation, with funding from the Federal German Ministry for Education and Research and by the European Research Council and the Israel Science Foundation.

## Supplementary Data

**Supplementary figure 1.**
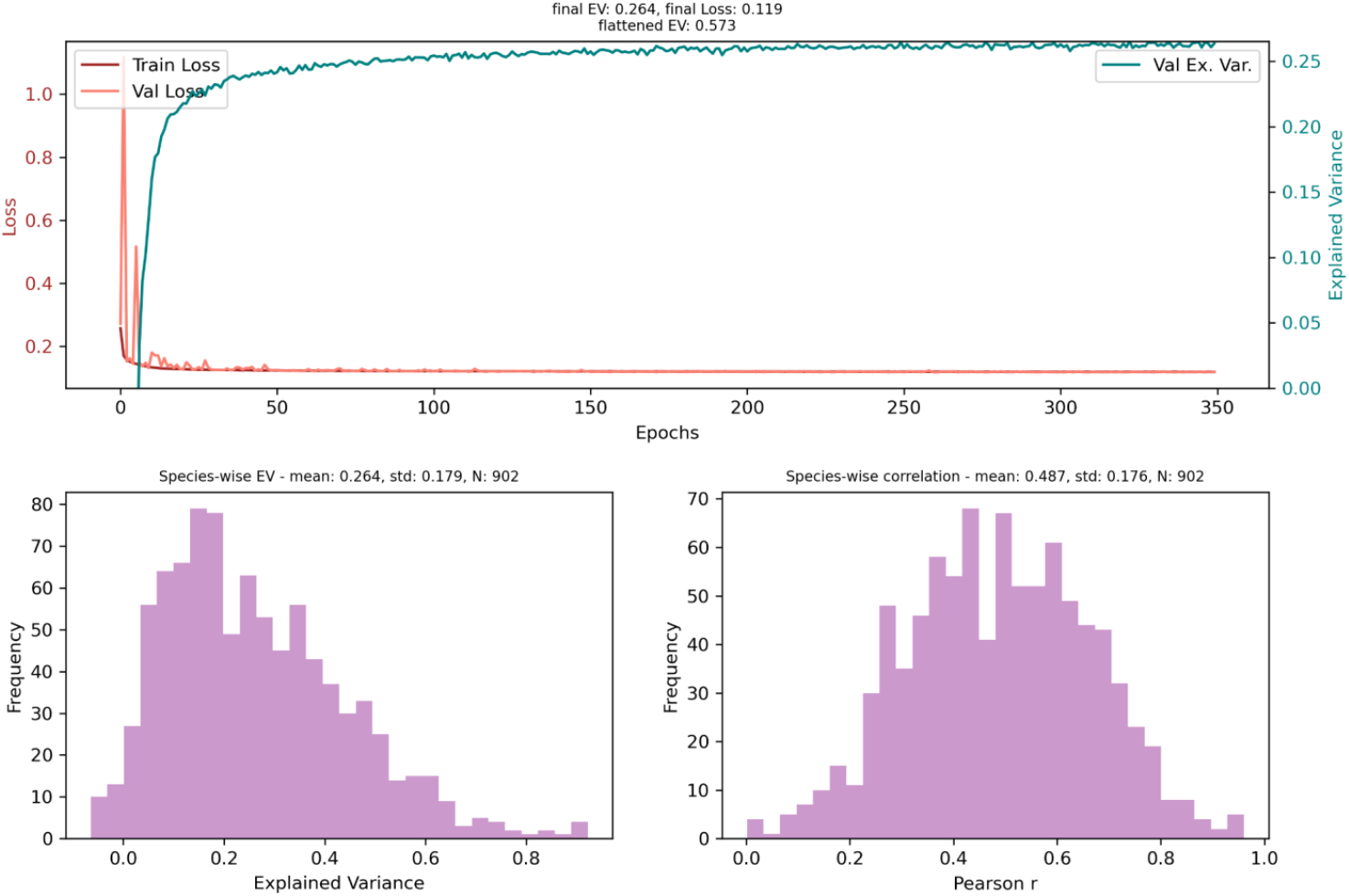
performances of the MAE-10% model reconstructing masked abundances. Top: train loss, validation loss, and explained variance on the validation set per epoch. Explained variance was calculated per species, and then averaged. Bottom: distribution of the average explained (left) or correlation between original and reconstructed abundances (right) of each species, calculated on the validation set in the final epoch.

**Supplementary figure 2.**
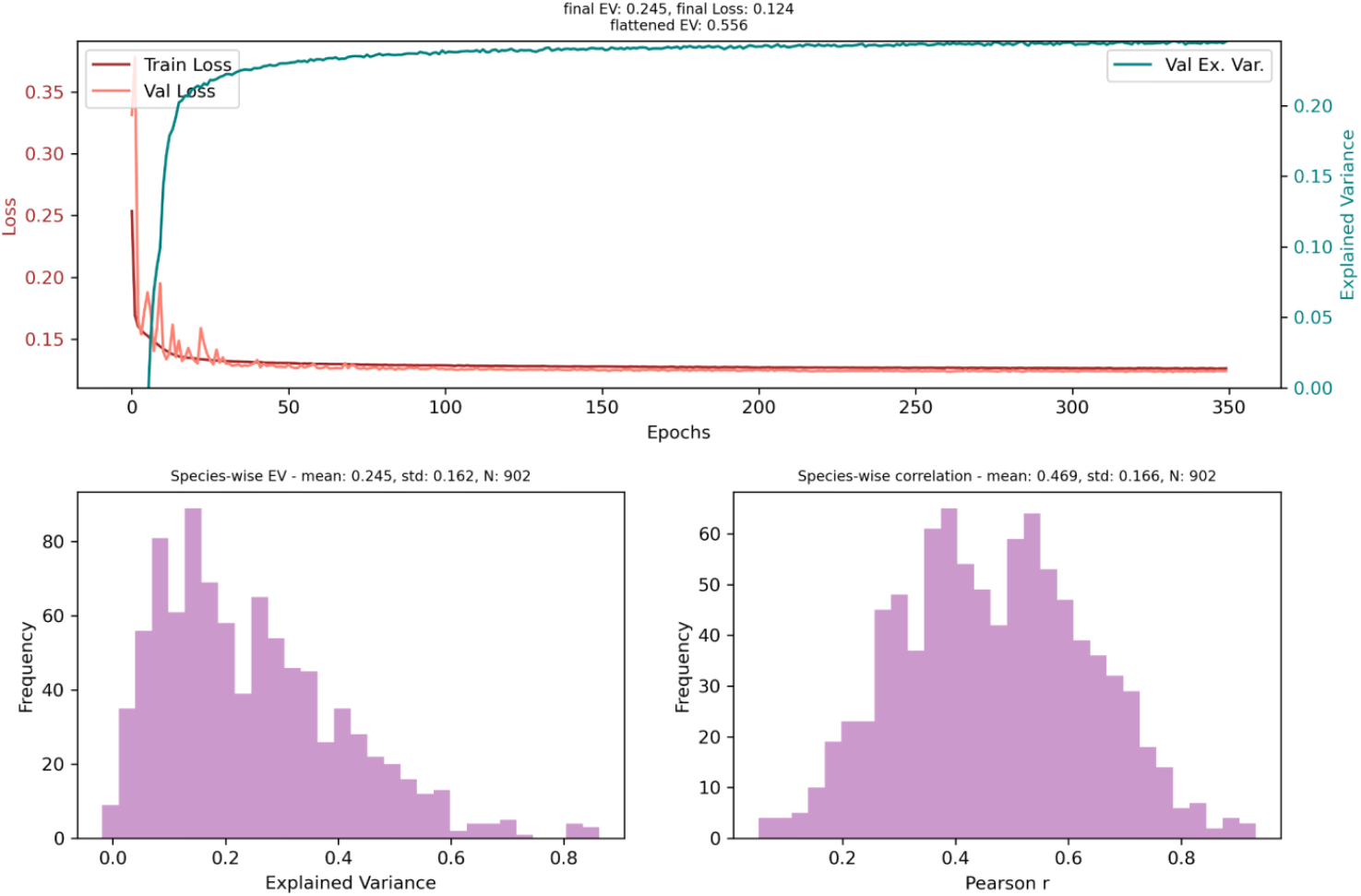
performances of the MAE-30% model reconstructing masked abundances. Top: train loss, validation loss, and explained variance on the validation set per epoch. Explained variance was calculated per species, and then averaged. Bottom: distribution of the average explained variance (left) or correlation between original and reconstructed abundances (right) of each species, calculated on the validation set in the final epoch.

**Supplementary figure 3.**
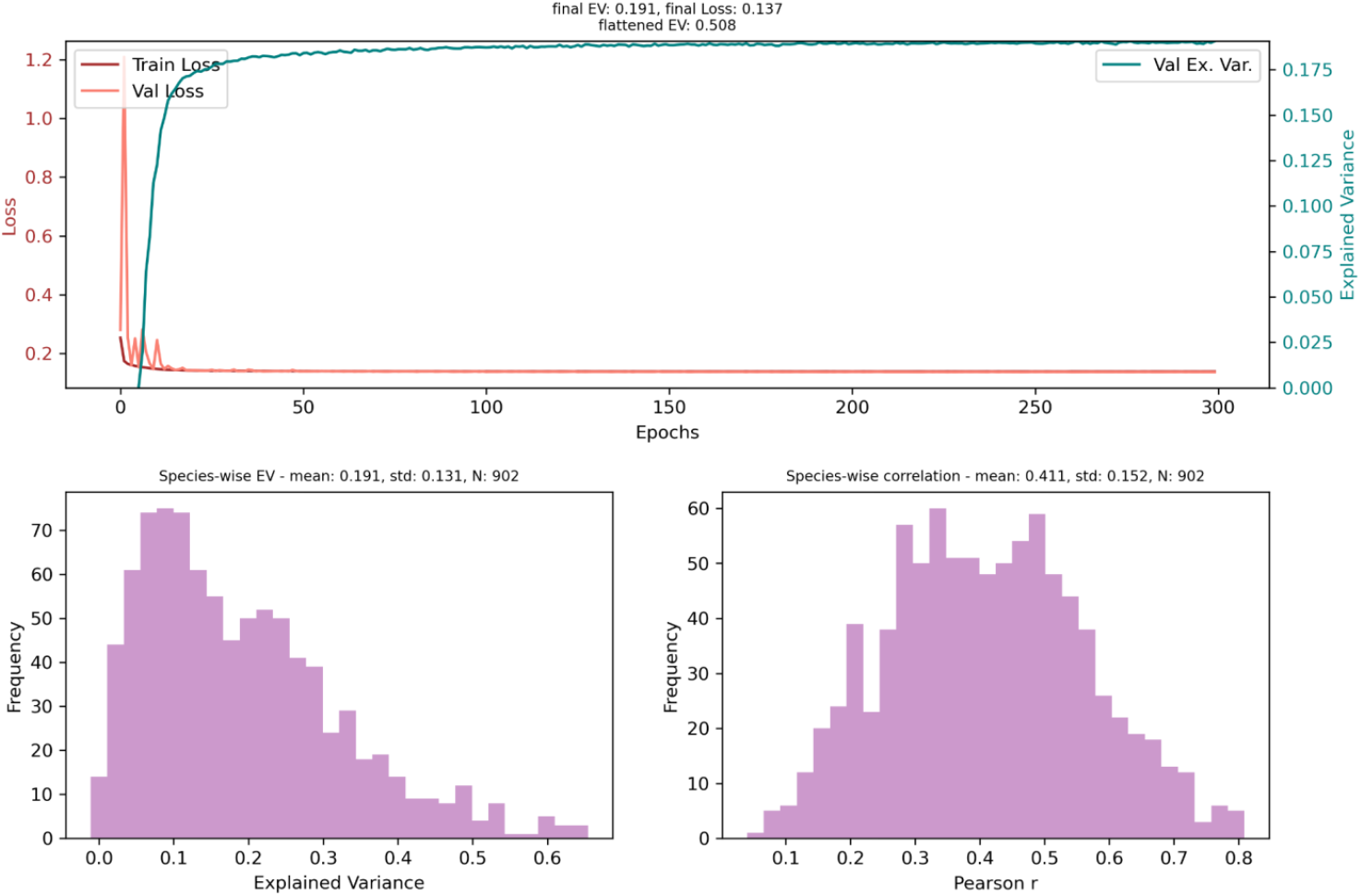
performances of the MAE-70% model reconstructing masked abundances. Top: train loss, validation loss, and explained variance on the validation set per epoch. Explained variance was calculated per species, and then averaged. Bottom: distribution of the average explained variance (left) or correlation between original and reconstructed abundances (right) of each species, calculated on the validation set in the final epoch.

**Supplementary table 1.**
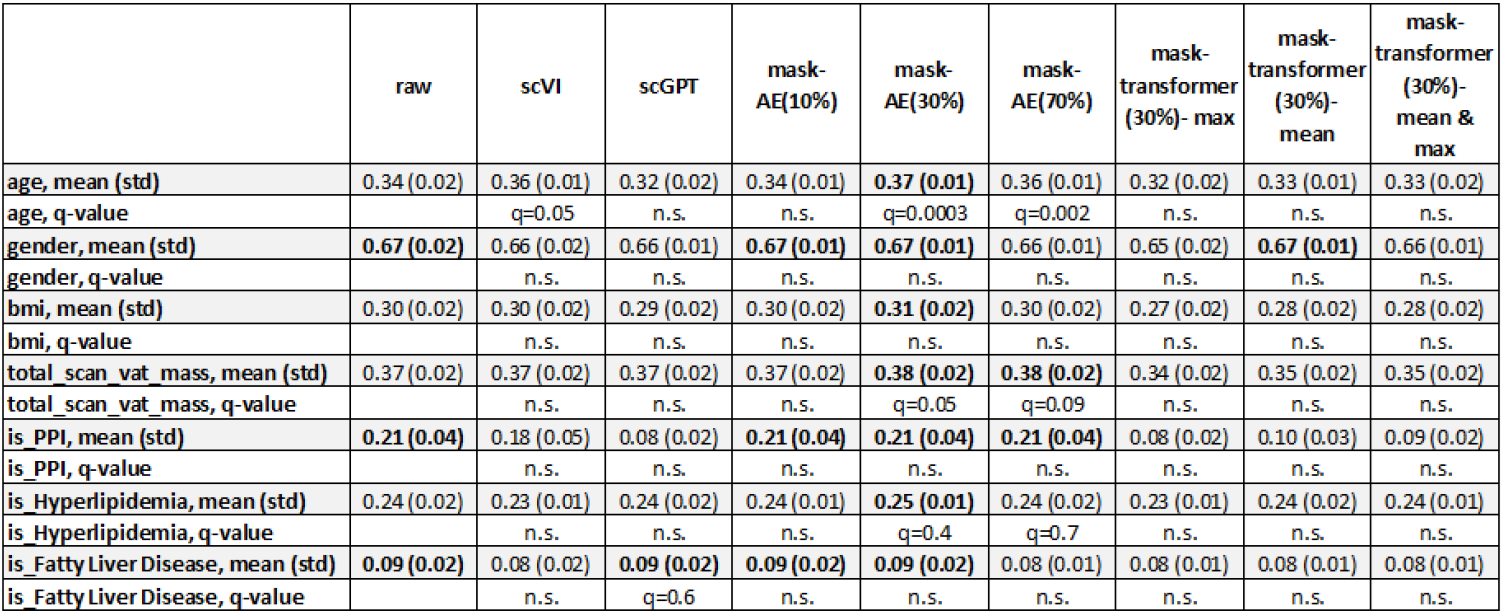
Prediction performance across different microbiome representations, including multiple pooling strategies. Performance evaluation was based on four repetitions of a 5-fold cross-validation, resulting in a total of 20 train-test split experiments. The table shows the performance (continuous: Pearson correlation, binary: PR-AUC) mean and standard deviation across all test sets. Q-values are based on the Mann-Whitney U test comparing each performance distribution to the raw model’s performances, and corrected for multiple testing using FDR.

|  | raw | scVI | scGPT | mask-AE(10%) | mask-AE(30%) | mask-AE(70%) | mask-transformer (30%)- max | mask-transformer (30%)- mean | mask-transformer (30%)- mean & max |
| --- | --- | --- | --- | --- | --- | --- | --- | --- | --- |
| age, mean (std) | 0.34 (0.02) | 0.36 (0.01) | 0.32 (0.02) | 0.34 (0.01) | **0.37 (0.01)** | 0.36 (0.01) | 0.32 (0.02) | 0.33 (0.01) | 0.33 (0.02) |
| age, q-value |  | q=0.05 | n.s. | n.s. | q=0.0003 | q=0.002 | n.s. | n.s. | n.s. |
| gender, mean (std) | **0.67 (0.02)** | 0.66 (0.02) | 0.66 (0.01) | **0.67 (0.01)** | **0.67 (0.01)** | 0.66 (0.01) | 0.65 (0.02) | **0.67 (0.01)** | 0.66 (0.01) |
| gender, q-value |  | n.s. | n.s. | n.s. | n.s. | n.s. | n.s. | n.s. | n.s. |
| bmi, mean (std) | 0.30 (0.02) | 0.30 (0.02) | 0.29 (0.02) | 0.30 (0.02) | **0.31 (0.02)** | 0.30 (0.02) | 0.27 (0.02) | 0.28 (0.02) | 0.28 (0.02) |
| bmi, q-value |  | n.s. | n.s. | n.s. | n.s. | n.s. | n.s. | n.s. | n.s. |
| total_scan_vat_mass, mean (std) | 0.37 (0.02) | 0.37 (0.02) | 0.37 (0.02) | 0.37 (0.02) | **0.38 (0.02)** | **0.38 (0.02)** | 0.34 (0.02) | 0.35 (0.02) | 0.35 (0.02) |
| total_scan_vat_mass, q-value |  | n.s. | n.s. | n.s. | q=0.05 | q=0.09 | n.s. | n.s. | n.s. |
| is_PPI, mean (std) | **0.21 (0.04)** | 0.18 (0.05) | 0.08 (0.02) | **0.21 (0.04)** | **0.21 (0.04)** | **0.21 (0.04)** | 0.08 (0.02) | 0.10 (0.03) | 0.09 (0.02) |
| is_PPI, q-value |  | n.s. | n.s. | n.s. | n.s. | n.s. | n.s. | n.s. | n.s. |
| is_Hyperlipidemia, mean (std) | 0.24 (0.02) | 0.23 (0.01) | 0.24 (0.02) | 0.24 (0.01) | **0.25 (0.01)** | 0.24 (0.02) | 0.23 (0.01) | 0.24 (0.02) | 0.24 (0.01) |
| is_Hyperlipidemia, q-value |  | n.s. | n.s. | n.s. | q=0.4 | q=0.7 | n.s. | n.s. | n.s. |
| is_FattyLiverDisease, mean (std) | **0.09 (0.02)** | 0.08 (0.02) | **0.09 (0.02)** | **0.09 (0.02)** | **0.09 (0.02)** | 0.08 (0.01) | 0.08 (0.01) | 0.08 (0.01) | 0.08 (0.01) |
| is_FattyLiverDisease, q-value |  | n.s. | q=0.6 | n.s. | n.s. | n.s. | n.s. | n.s. | n.s. |

**Supplementary table 2.** Prediction performance across microbiome representations with limited (n=100) training data, including multiple pooling strategies. Train cohorts consisted of 100 randomly selected samples. Performance distribution is based on 100 repetitions. The table shows the performance (continuous: Pearson correlation, binary: PR-AUC) mean and standard deviation across all test sets. Q-values are based on the Mann-Whitney U test comparing each performance distribution to the raw model’s performances, and corrected for multiple testing using FDR.

|  | raw | scVI | scGPT | mask-AE(10%) | mask-AE(30%) | mask-AE(70%) | mask-transformer (30%)- max | mask-transformer (30%)- mean | mask-transformer (30%)- mean & max |
| --- | --- | --- | --- | --- | --- | --- | --- | --- | --- |
| age, mean (std) | 0.06 (0.11) | **0.14 (0.11)** | 0.13 (0.09) | 0.10 (0.11) | 0.11 (0.11) | 0.11 (0.12) | 0.11 (0.09) | 0.12 (0.11) | 0.12 (0.10) |
| age, q-value |  | q<10<sup>-5</sup> | q<10<sup>-4</sup> | q=0.08 | q=0.003 | q=0.004 | q=0.004 | q=0.003 | q=0.002 |
| gender, mean (std) | 0.62 (0.07) | 0.60 (0.07) | 0.63 (0.06) | 0.62 (0.06) | **0.64 (0.06)** | **0.64 (0.06)** | 0.61 (0.06) | 0.61 (0.06) | 0.61 (0.06) |
| gender, q-value |  | n.s. | q=0.2 | q=0.7 | q=0.2 | q=0.08 | n.s. | n.s. | n.s. |
| bmi, mean (std) | 0.11 (0.11) | 0.12 (0.11) | 0.15 (0.09) | 0.15 (0.11) | **0.16 (0.11)** | **0.16 (0.11)** | 0.11 (0.11) | 0.13 (0.11) | 0.12 (0.11) |
| bmi, q-value |  | q=0.8 | q=0.03 | q=0.08 | q=0.003 | q=0.02 | q=0.8 | q=0.5 | q=0.7 |
| total_scan_vat_mass, mean (std) | 0.18 (0.11) | 0.18 (0.10) | 0.20 (0.10) | 0.23 (0.10) | **0.25 (0.11)** | 0.24 (0.11) | 0.17 (0.11) | 0.17 (0.11) | 0.18 (0.11) |
| total_scan_vat_mass, q-value |  | q=0.9 | q=0.4 | q=0.004 | q=0.0004 | q=0.003 | q=0.9 | q=0.9 | q=0.8 |
| is_PPI, mean (std) | 0.73 (0.08) | 0.76 (0.06) | 0.61 (0.06) | 0.79 (0.06) | 0.80 (0.05) | **0.81 (0.04)** | 0.60 (0.06) | 0.63 (0.06) | 0.62 (0.06) |
| is_PPI, q-value |  | q=0.009 | n.s. | q<10<sup>-5</sup> | q<10<sup>-9</sup> | q<10<sup>-11</sup> | n.s. | n.s. | n.s. |
| is_Hyperlipidemia, mean (std) | **0.53 (0.05)** | **0.53 (0.05)** | **0.53 (0.05)** | 0.52 (0.06) | 0.52 (0.05) | 0.52 (0.05) | 0.52 (0.05) | **0.53 (0.06)** | **0.53 (0.05)** |
| is_Hyperlipidemia, q-value |  | q=0.6 | q=0.8 | q=0.9 | q=0.9 | q=0.8 | q=0.8 | q=0.7 | q=0.7 |
| is_FattyLiverDisease, mean (std) | 0.57 (0.05) | 0.55 (0.06) | 0.58 (0.06) | 0.58 (0.06) | **0.59 (0.07)** | 0.58 (0.06) | 0.57 (0.06) | 0.57 (0.06) | 0.57 (0.06) |
| is_FattyLiverDisease, q-value |  | n.s. | q=0.3 | q=0.4 | q=0.1 | q=0.5 | q=0.9 | q=0.7 | q=0.8 |

**Supplementary figure 4.**
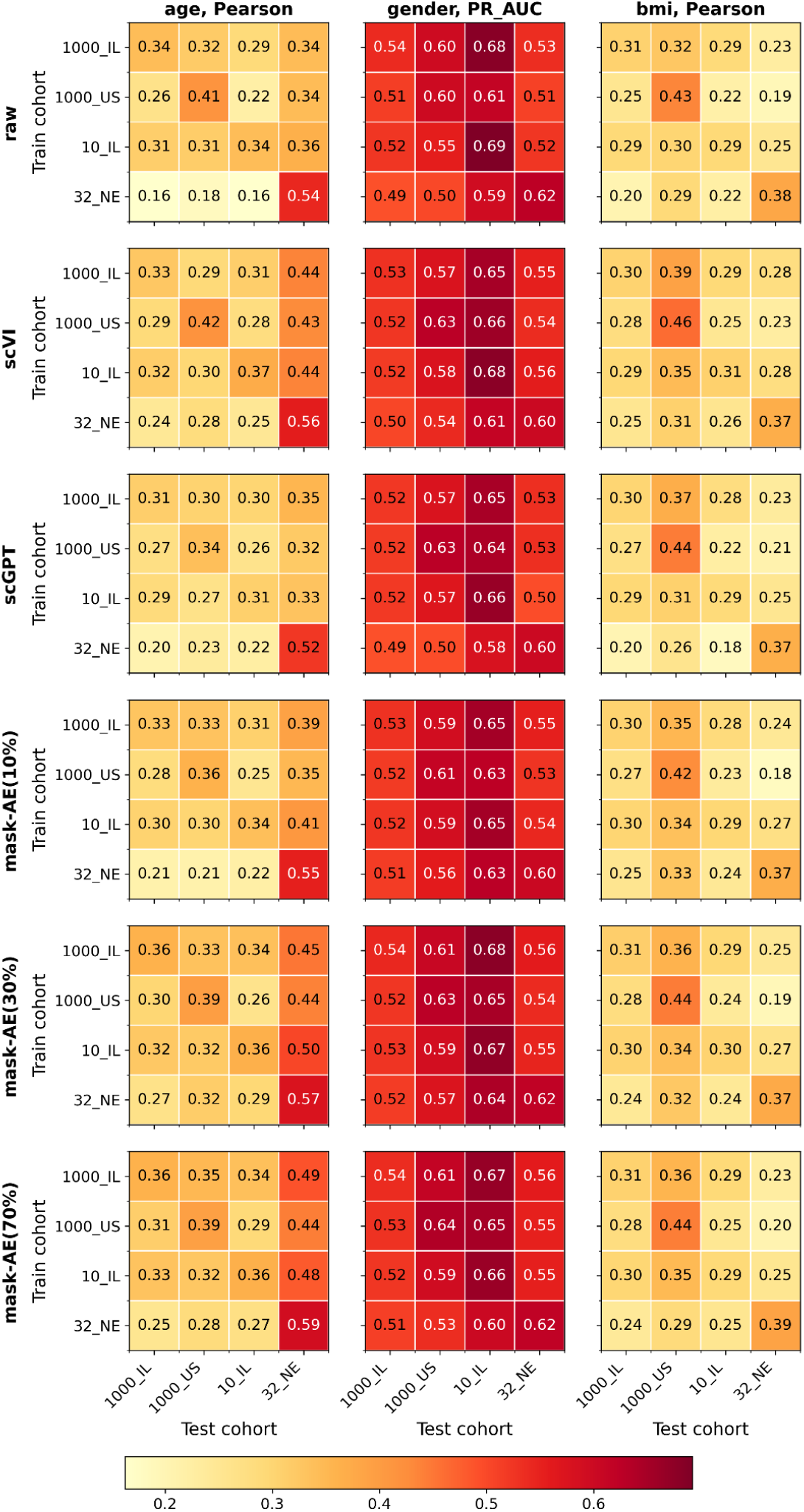
test performance of supervised predictors trained on one cohort and tested on another.

## References

1. Jumper, J. et al. Highly accurate protein structure prediction with AlphaFold. Nature 596, 583–589 (2021).

2. Ji, Y., Zhou, Z., Liu, H. & Davuluri, R. V. DNABERT: pre-trained Bidirectional Encoder Representations from Transformers model for DNA-language in genome. Bioinformatics 37, 2112– 2120 (2021).

3. Lopez, R., Regier, J., Cole, M. B., Jordan, M. I. & Yosef, N. Deep generative modeling for single-cell transcriptomics. Nat. Methods 15, 1053–1058 (2018).

4. Theodoris, C. V. et al. Transfer learning enables predictions in network biology. Nature 618, 616– 624 (2023).

5. Pope, Q., Varma, R., Tataru, C., David, M. & Fern, X. Learning a deep language model for microbiomes: the power of large scale unlabeled microbiome data. BioRxiv (2023) doi:10.1101/2023.07.17.549267.

6. Zhang, H. et al. MGM as a large-scale pretrained foundation model for microbiome analyses in diverse contexts. BioRxiv (2025) doi:10.1101/2024.12.30.630825.

7. Oh, M. & Zhang, L. DeepGeni: deep generalized interpretable autoencoder elucidates gut microbiota for better cancer immunotherapy. Sci. Rep. 13, 4599 (2023).

8. Oh, M. & Zhang, L. DeepMicro: deep representation learning for disease prediction based on microbiome data. Sci. Rep. 10, 6026 (2020).

9. Boiarsky, R. et al. Deeper evaluation of a single-cell foundation model. Nat. Mach. Intell. 6, 1443– 1446 (2024).

10. Wong, D. R., Hill, A. S. & Moccia, R. Simple controls exceed best deep learning algorithms and reveal foundation model effectiveness for predicting genetic perturbations. Bioinformatics 41, (2025).

11. Rothschild, D. et al. An atlas of robust microbiome associations with phenotypic traits based on large-scale cohorts from two continents. PLoS ONE 17, e0265756 (2022).

12. Zahavi, L. et al. Bacterial SNPs in the human gut microbiome associate with host BMI. Nat. Med. 29, 2785–2792 (2023).

13. Reicher, L. et al. Deep phenotyping of health-disease continuum in the Human Phenotype Project. Nat. Med. (2025) doi:10.1038/s41591-025-03790-9.

14. Gacesa, R. et al. Environmental factors shaping the gut microbiome in a Dutch population. Nature 604, 732–739 (2022).

15. Cui, H. et al. scGPT: toward building a foundation model for single-cell multi-omics using generative AI. Nat. Methods 21, 1470–1480 (2024).

16. Grinsztajn, L., Oyallon, E. & Varoquaux, G. Why do tree-based models still outperform deep learning on typical tabular data? Advances in Neural Information Processing Systems (2022).

17. Leviatan, S., Shoer, S., Rothschild, D., Gorodetski, M. & Segal, E. An expanded reference map of the human gut microbiome reveals hundreds of previously unknown species. Nat. Commun. 13, 3863 (2022).

18. Ke, G. et al. LightGBM: A Highly Efficient Gradient Boosting Decision Tree. Advances in Neural Information Processing Systems (2017).

19. Virtanen, P. et al. SciPy 1.0: fundamental algorithms for scientific computing in Python. Nat. Methods 17, 261–272 (2020).

20. Benjamini, Y. & Yekutieli, D. The Control of the False Discovery Rate in Multiple Testing under Dependency. Annals of statistics (2001).

